# Comprehensive network analysis reveals alternative splicing-related lncRNAs in hepatocellular carcinoma

**DOI:** 10.1101/548263

**Authors:** Junqing Wang, Yixin Chen, Keli Xu, Yin-yuan Mo, Yunyun Zhou

**Affiliations:** Department of Surgery, Ruijin Hospital, Shanghai Jiao Tong University School of Medicine, Shanghai, 20025, China; Department of Computer and Information Science, University of Mississippi, Oxford, MS, 38655; Department of Neurobiology and Anatomical Sciences, University of Mississippi Medical Center, Jackson, Mississippi, 39216; Department of Pharmacology and Toxicology, University of Mississippi Medical Center, Jackson, Mississippi, 39216; Department of Data Science, University of Mississippi Medical Center, Jackson, Mississippi, 39216, USA

**Keywords:** lncRNA, alternative splicing, random walk, network analysis

## Abstract

A number of recent studies have highlighted the findings that certain lncRNAs are associated with alternative splicing (AS) in tumorigenesis and progression. Although existing work showed the importance of linking certain misregulations of RNA splicing with lncRNAs, a primary concern is the lack of genome-wide comprehensive analysis for their associations.

We analyzed an extensive collection of RNA-seq data, quantified 198,619 isoform expressions, and found systematic isoform usage changes between hepatocellular carcinoma (HCC) and normal liver tissue. We identified a total of 1375 splicing switched isoforms and further analyzed their biological functions.

To predict which lncRNAs are associated with these AS genes, we integrated the co-expression networks and epigenetic interaction networks collected from text mining and database searching, linking lncRNA modulators such as splicing factors, transcript factors, and miRNAs with their targeted AS genes in HCC. To model the heterogeneous networks in a single framework, we developed a multi-graphic random walk (RWMG) network method to prioritize the lncRNAs associated with AS in HCC. RWMG showed a good performace evaluated by ROC curve based on cross-validation and bootstrapping strategy.

As a summary, we identified 31 AS-related lncRNAs including MALAT1 and HOXA11-AS, which have been reported before, as well as some novel lncRNAs such as DNM1P35, HAND2-AS1, and DLX6-AS1. Survival analysis further confirmed the clinical significance of identified lncRNAs.

## 1. Introduction

Alternative splicing (AS) changes are frequently observed in cancer and may serve as the cancer driver genes. They could originate from somatic mutations that dysfunctions the splicing regulatory mechanisms or influences the expression changes of splicing factors or transcript factors^1^. Therefore, AS genes recognized as important signatures for tumorigenesis are of significant values in developing therapeutic targets in cancer clinical trials. For example, SF3B1-targeting compounds spliceosome inhibitor E7107 which have been implemented in advanced tumor^2^.

Numerous studies showed that the long non-coding RNAs “lncRNAs” (>200nt in length) are associated with a number of AS mechanisms^3,4^. LncRNAs may interact with specific alternative splicing factors (ASF) or through other intermediate molecules affecting chromatin remodeling to fine-tune the splicing of target genes^4^. For instance, our previous experimental study showed the MALAT1 regulated a ASF, SRSF1 (SF2), in gastric cancer cells^5,6^. Ji et al. reported MALAT1 promotes tumor growth and metastasis in colorectal cancer through binding to SFPQ and releasing oncogene PTBP2 ^7^. LINC01133 interacts with splicing factor SRSF6 in colorectal cancer ^8^, acting as a target mimic for SRSF6 interaction with EZH2 and LSD1 in non-small cell lung cancer (NSCLC)^9^.

A number of splicing regulatory proteins that promote the transformation of their target genes can be triggered by transcriptional factors (TFs). For example, a TF, MYC, induced upregulation of hnRNP A1 and hnRNP A2, which in turn, regulate alternative splicing of pyruvate kinase to promote expression of the cancer-associated pyruvate kinase M2 (PKM2) isoform^10,11^. There exist more comprehensive lncRNA regulatory mechanism in AS, since lncRNAs are either pre-transcriptional or post-transcriptional specialists, acting as decoys to draw effectors (miRNAs, TFs, or ASFs) away from their targets, as cofactors or guides to alter TF-promoter interactions, and as molecular switches to alter TF or ASF activity across multiple targets.

Although previous efforts identified both lncRNAs and AS that may be important in cancer, gaps exist in current studies that only a few cancer-related AS that are regulated by lncRNAs have been extensively investigated. However, their methodology either didn’t genome-widely link lncRNAs to comprehensive AS mechanisms or correlate with clinical outcomes. Furthermore, to date, next-generation sequencing technologies have facilitated the identification of an accumulation of ∼40K novel lncRNAs, whose regulatory functions in AS remain unknown in tumorigeneses. Therefore, a key open question is: how many novel lncRNAs are associated with AS modulations in tumorigenesis, genome-wisely.

Here we utilized a novel network propagation technology, random walk-based multi-graphic model (RWMG), to simultaneously integrate complicated biological connections among lncRNA -effectors (TF, ASF, and miRNA) -AS interaction networks and co-expression networks in a single analysis framework. This method is an extended application inspired by Random walk with restart algorithm to prioritize important lncRNAs that are involved in AS based on the hypothesis that more important genes are likely to receive more links from other networks. In comparison with traditional random walk algorithms, which treat all genes equally, our flexible, scalable method can be formulated to rank a subset of vertices (e.g., PCGs, lncRNAs), based on pre-knowledge as the starting walking vertices. This method is more accurate than other traditional “shortest path” network-based integrative methods, as it can overcome the “noisy” and “incomplete” highly dimensional heterogeneous data.

In addition, previous tumor and normal comparison studies are limited to normal adjacent to tumor (NAT) tissues. However, these tissues are not truly ‘normal’ as they usually surrounded by tumor contaminations. Therefore, many potential cancer biomarkers involved in AS may be missed. By combining the ‘pure’ healthy liver organs from GTEx with TCGA expression data, we broaden the scope of suspected candidates with a variety of AS patterns.

## 2. Method and materials

### 2.1 Data description and project design

The framework of the underlying biological hypothesis and model assumption for this project can be found in Fig S1 A-B. The analysis described in this manuscript relied on multiple types of data. We downloaded GTEx RNAseq data with 110 normal liver samples; TCGA RNAseq data and clinical data from 369 liver tumors and 50 normal samples from the UCSC Xena database (http://xena.ucsc.edu/). The raw RNAseq data (Illumina HiSeq 2000 RNA Sequencing platform) was re-processed with UCSC’s Xena Toil ^12^ to quantify gene level and transcript isoform level expression for both coding and non-coding genes. Given that the non-coding transcript expression of RNAseq data contains many small and uncertain transcripts, we filtered out the small and uncertain transcripts but kept transcripts for intergenic lncRNAs, antisense, sense_intronic, sense_overlapping, processed_transcripted, and processed_pseudogene categories based on GENCODE v23 annotation^13^.

### 2.2 Identification of HCC Tumor-specific non-coding genes (lncRNAs, pseudogenes) from TCGA and GTEx RNA-seq data

We performed a trimmed mean of M-values (TMM) normalization method for RNAseq count data ^14^ so that the expression level for lncRNAs and pseudogenes are comparable. The TMM normalized data were further transformed to log2-counts per million for linear modeling. HCC differentially expressed (DE) lncRNAs and pseudogenes between tumor and normal samples (T/N) were analyzed by R package limma^15^ with cutoff settings (P<1.0E-04 and Fold Change > 2). The method to identify DE miRNAs has been reported in our previous work^16^. These identified HCC specific non-coding DE genes (lncRNAs, pseudogenes, and miRNAs) are expected to represent potential key functions in liver tumorigenesis.

### 2.3 Analysis of alternative splicing isoforms and functional consequence

We first removed isoforms which are all zero counts across all the samples. We used R package “IsoformSwitchAnalyzeR” to analyze individual isoform switches from T/N comparison and their biological consequence changes^17,18^. Differentially switched isoforms between T/N were determined by the following criteria: difference in isoform fraction (dIF) > 0.1 and FDR corrected q-value < 0.05. The functional consequences of switched isoforms were further analyzed for protein-coding potential(CPAT)^19^, Non-sense mediated decay (NMD) status, protein domains (Pfam) ^20,21^, and the amino acid sequence of open reading frames (ORF). We used 0.364 as suggested to distinguish coding and non-coding isoforms in CPAT analysis. NMD is a process that recognizes mRNAs carrying a premature termination codon (PTC) and triggers their degradation to prevent the synthesis of dysfunctional or even harmful proteins. AS that controls gene expression is an important process facilitating mRNA degradation in specific isoforms that would lead to NMD ^22^. Since we know the exon structure of all isoforms in a given gene (with isoform switching), we obtained their corresponding spliced nucleotide sequence and corresponding coding sequence from ORF positions^23^. The alternative splicing (AS) patterns of switched isoforms predicted by spliceR^24^ include Alternative 3’ acceptor sites (A3), Alternative 5’ donor sites (A5), Exon skipping (ES), Mutually exclusive exons (MEE), Mutually exclusive skip (MES), AS at TF start sites (ATSS), AS at termination site (ATTS), and Intron retention (IR). Enrichment tests are performed via base R’s prop.test and comparisons of group difference are done with fisher.test. P values are corrected for multiple testing using the Benjamin-Hochberg scheme and an FDR < 0.05 is considered significant.

### 2.4 Construction of AS-associated lncRNA epigenetic regulatory interaction subnetworks in HCC

We collected all possible physical interactions of lncRNAs and their targeted genes through database searching and text mining as the global background information. These interactions are evidenced from experiments validation, neighborhood, gene fusion, and co-occurrence information of lncRNAs connecting with miRNA-, TF-, ASF-, and switched genes. Specifically, HCC LncRNA-target networks were compiled from the following resources: Chiu et al.,^25^, miWalker2.0 ^26^, STARBASE v2 ^27^, and lncRNA-disease ^28 29^, which were analyzed from several high-throughput assays, including ENCODE enhanced version of the crosslinking and immunoprecipitation assay (eCLIP) and chromatin immunoprecipitation sequencing (ChIP-seq) data ^30^. HCC specific miRNA-target networks has been described in our previous published results^16^; TF-target predicted interaction network were manually curation from the following databases and publications Chiu et al. ^25, Table S5^, HTRIdb ^31^, Whitfield ^32^, and TRANSFAC ^33^ based on combined evidence from ENCODE ChIP-Seq assays and position weighted matrix (PWM) for TF motif analysis.

Genes related to AS regulatory pathway were collected from pathCards^34^, KEGG spliceosome ^35^, NCBI Biosystems mRNA processing^36^, REATOME mRNA splicing pathway and processing of capped intron-containing pre-mRNA pathway^37^. These genes were involved in an essential component of splicing factors or non-snRNA spliceosome required for the second catalytic step of pre-mRNA splicing. Among these collected 335 splicing regulator genes, 86 are experimental validated as alternative splicing factors (ASF). ASF and target genes interactions were manually confirmed from SpliceAid 2^38^, ASF motif analysis from SFmap^39^, a subset of RNA-binding protein network by Chiu et. al^25, Table S6^, and STRING database ^40^.

Finally, identified HCC DE lncRNAs, pseudogenes, and miRNAs were mapped to the global regulatory networks to construct HCC-specific sub-networks that contain switched genes as the targets or TF/ASF as the co-effectors of non-coding RNA regulators.

### 2.5 Construction of HCC lncRNA-AS co-expression interaction networks at isoform level

Pearson correlation was used to estimate the lncRNA co-expression relationships at isoform level. We only kept the connections for the pairs of lncRNA and protein-coding genes (PCGs), if their absolute correlation coefficient > 0.75, FDR p <0.05. The types of PCGs included the TFs, ASFs, and genes with isoform switches. Directions of lncRNAs that are negatively correlated with their targeted PCGs were predicted to inhibit their expression, while positive correlation indicates activation.

### 2.6 Random walk multi-graphic (RWMG) model for the integration of heterogeneous interaction networks

Random walk multi-graphic (RWMG) model is an integrative application of page rank with restart algorithm (RWR) on multiple layers of networks. Detailed method description can be found in our previously published report ^41^. Briefly, given a graph *G(V,E)*, (*i,j*)∈*V* are the vertices lncRNA *i* and AS genes *j,* and edge (*i,j*)∈*E* is weighted by the connectivity score between these vertices. Multiple edges are allowed to connect between any two vertices based on the relationship defined from the co-expression network, epigenetic regulatory network and splicing pathway PPI networks. Assuming the total number of lncRNAs to be *m* and of AS genes to be *n*, the probability for which lncRNA *i* will traverse to AS genes *j* is defined by the adjacency matrix: 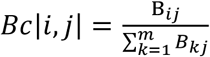. A “teleportation term” is added to *B*_*c*_ for the sake of numerical stability. Thus the transition matrix for networks is defined as 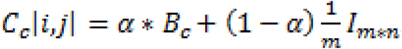, where *I*_*m*n*_ *is* an *m* by *n* matrix with all entries is equal to 1, α is the probability of a lncRNA jumping to one of its neighbors with the probability governed by matrix *Bc*, and 1–*α* defines the probability of this lncRNA jumping randomly to any other vertex and relinquishing the matrix *Bc* for traversal.

The final multi-dimensional heterogeneous network will be merged for the overlapped lncRNA node features and taken the union of distinct nodes to augment each individual network with missing connections. We implemented the RWR algorithm on the final multi-graphic network by R packages dnet and igraph ^42^. Network visualization was performed by R package visNetwork ^43^. Those genes with known roles in regulating AS network will be set as the “seed” nodes in advance to predict the “new” lncRNAs, based on move probabilities from the current node to any of their randomly selected neighbors.

To evaluate our approach’s sensitivity, we simulated different random walk strategies for optimization. We created a list of experimentally validated AS associated genes as “gold-standard” true positive genes (TPG) curated from the careful literature review and randomly selected genes as the “gold-standard” true negative (TNG). We chose the “best” model that has the most candidate significantly enriched in the “gold-standard” gene list. In reality, the number of TPG is much smaller compared to TNG. To avoid bias from highly imbalanced data between these two sets, we performed a bootstrap resampling technique by selecting an equal number of data as TNG. This process was repeated 10 times, and the overall performances were calculated by the mean value of these performances.

### 2.7 Survival analysis for prognostic confirmation of identified pathogenic lncRNAs and Pseudogenes

To confirm the pathogenic characters of identified lncRNAs and pseudogenes, univariate Cox proportional model was used to evaluate the association of selected genes with overall survival outcomes. Kaplan–Meier plots and log-rank test statistics were used to visualize the high-and low-risk groups. The cutoff of the high-and low-risk group was determined by the median value of the normalized count of selected genes.

## 3. Results

### 3.1 Significantly expressed lncRNAs and pseudogenes in HCC

We identified 369 differentially expressed (DE) lncRNA genes and 171 DE pseudogenes from T/N comparison(Table S1). The visualizations of DE lncRNAs and pseudogenes were shown in volcano plots (Fig. 1). Compared to other studies, many DE LncRNAs, such as MALAT1, CDKN2B-AS1, and HOTTIP, have been reported to be associated with liver cancers before ^44-46^. We also highlighted several important pseudogenes, such as HNRNPA1P4, HNRNPA1P21, which are the pseudogenes of heterogeneous nuclear ribonucleoproteins A1 (hnRNPs) who play key roles in the regulation of alternative splicing. We performed DE analysis as the initial screen step to narrow the focus of the HCC specific non-coding genes associated with AS for the downstream network analysis.

**Figure 1.**
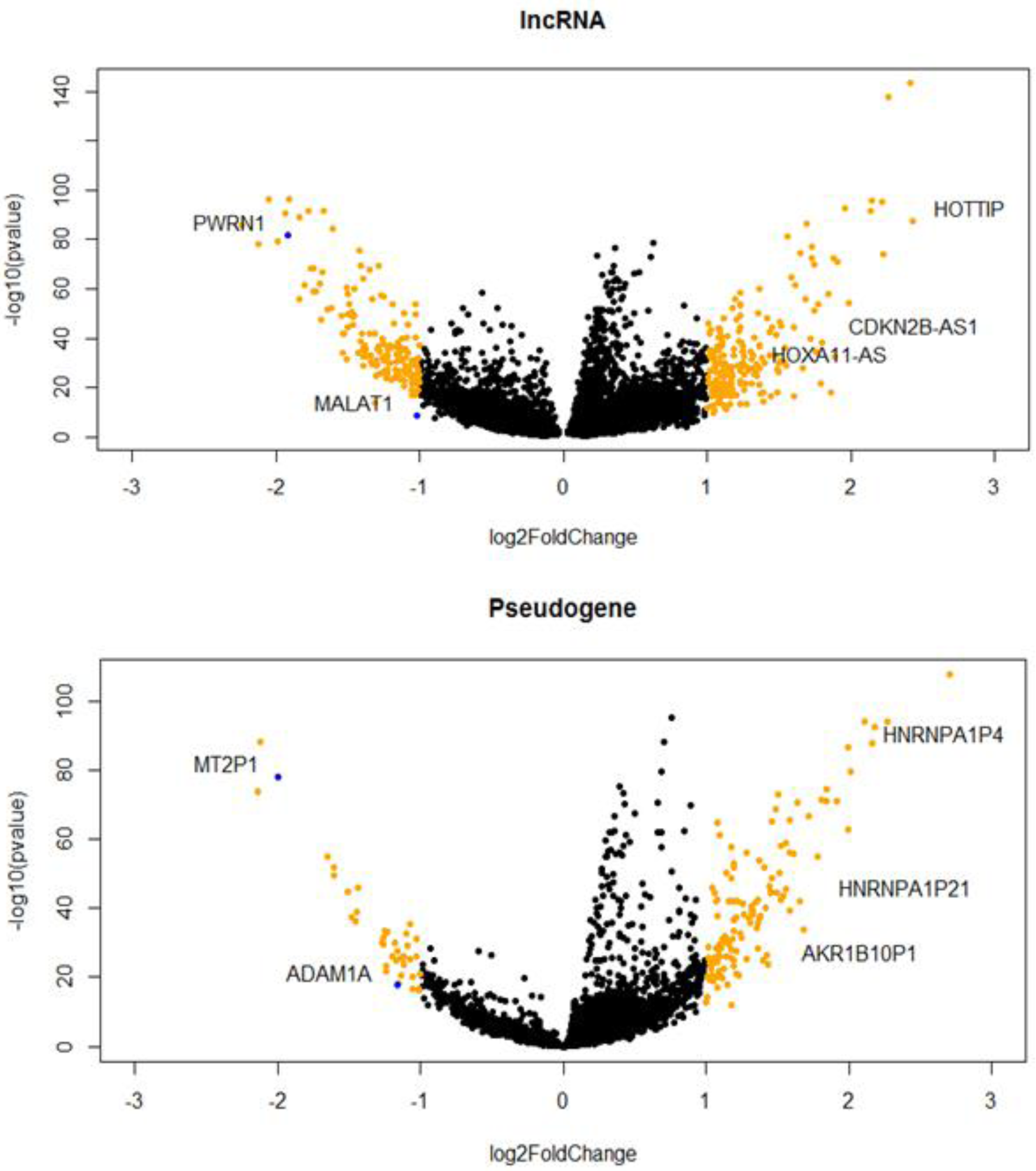
HCC specific lncRNAs (A) and Pseudogenes (B) that are differentially expressed in tumor and normal.

### 3.2 Identification of significantly switched isoforms and prediction of alternative splicing patterns

From the isoform level expression T/N comparison, we identified 1375 switched isoforms mapping to 1078 unique genes. Among these switched isoforms, 1251 are protein-coding isoforms, and 124 are non-coding isoforms including antisense, lincRNA, pseudogenes, and others (Table. S2). We found that the proportion of switching rate for coding genes is much higher than non-coding genes (Fisher’s exact test, p.value= 8.4e-08, Fig. 2 A-B). In order to intuitively visualize the splicing composition of these switched isoforms, we breakdown the dIF distribution according to isoform types such as lincRNA, antisense, and pseudogenes with the most significant switched isoforms (dIF > 0.2 or dIF < -0.2) highlighted (Fig. 2C).

**Figure 2.**
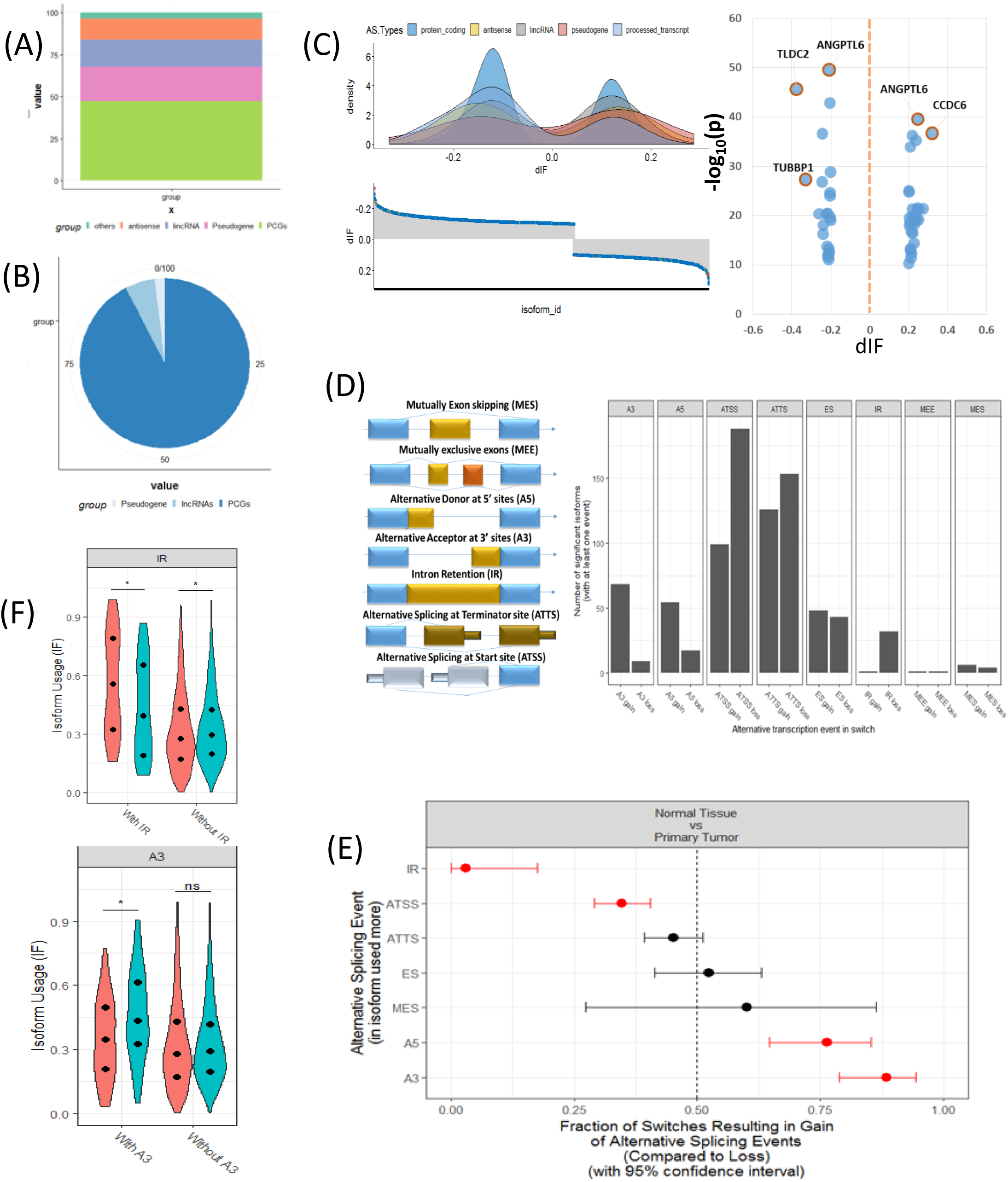
Genome-wide transcript analysis for switched isoforms between tumor and normal comparison in HCC. (A) Global distribution of whole genome transcriptions based on GENCODE annotation. The percentage of coding and non-coding genes are about half and half. (B) Distribution of the HCC switched Isoforms in coding and non-coding region. About 95% of switched isoforms are from protein-coding genes. (C) Distribution of differential isoform fraction (dIF) stratified by coding or non-coding isoform types. The most significantly switched Isoforms (dIF> 0.2) are highlighted. (D) Illustration of alternative splicing event types for the switched isoforms and distribution of Isoforms gain (increased dIF) or loss (decreased dIF) in each types. (E) Enrichment analysis for alternative splicing types in isoform fraction gain or loss. IR and ATSS categories are enriched in loss switches, while A5 and A3 are significantly enriched in gain. (F) Distribution of dIF changes with or without IR and A3 events. Isoforms showed less usages in IR type and more usage in A3 type.

Fig. 2D shows the eight splicing patterns for switched isoforms stratified by isoforms usage gain or loss in the tumor. Some of the switched isoforms are predicted to have multiple AS events in HCC (Table S3). Interestingly, we observed a global phenomenon that the AS events are not equally used -most prominently illustrated by the use of ATSS in HCC, where there is (a lot) more losses than the gain of amino acid coding exons. It is noteworthy that IR and ATSS are enriched in significant loss of isoform usage in tumor, but A5 and A3 are significantly enriched in a gain (Fig 2E). Here IR events are of particular functional interest since they represent the largest changes in isoforms. As we displayed in the violin plots, the enriched IR and A3 splicing groups reported the significantly opposite direction of isoform usages between T/N samples (Fig. 2F).

### 3.3 Analysis of functional consequences for switched isoforms

The overview of switched isoforms impacting the biological function alterations in HCC is shown in Fig 3A. We can see that the number of protein domain gain is comparable to domain loss, but is significantly more than domain “switch”. Here, the “switch” term indicates both a gain and a loss occurred. Also, switching resulting in ORF gain is significantly more than ORF loss. For the Gene Ontology analysis, both gain and loss switched isoforms are associated with different types of metabolic process. KEGG analysis shown the isoform loss in tumor tissue are associated with virus infection, hepatitis C, etc, while isoform gain in the tumor is associated with Base excision repair, apoptosis, etc (Table S2).

**Figure 3.**
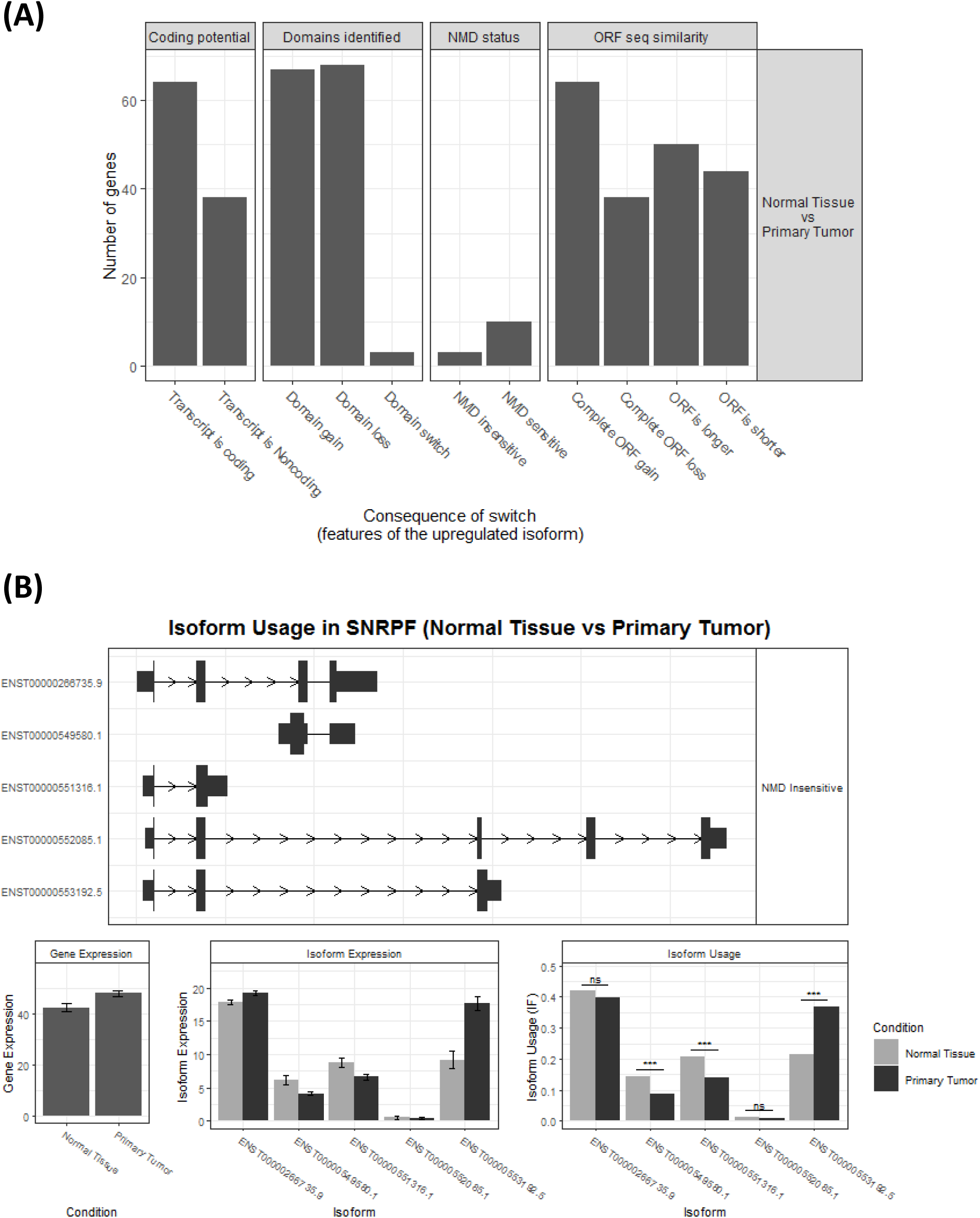
(A) Overview of the number of switched isoforms predicted to have functional consequences. (B) Visualization of switched Isoform structure. Taking an splicing factor gene, SNRPF, for example, its isoform ENST00000553192.5.1 showed oposite switching pattern compared to others. And 3 out of 5 isoforms showed differential isoform expressions, although no difference for the overall gene expression.

Importantly, we confirmed 20 genes with switched isoforms which are involved in AS regulatory functions (Table 1). Fig. 3B displays one of an example of AS factor, SNRPF’s isoforms structures, gene expression, and isoforms’ usage in T/N comparison. SNRPF is a core component of U small nuclear ribonucleoproteins, which are key components of the pre-mRNA processing spliceosome. We can see from the figure that there is no significant difference for SNRPF gene expression, but an opposite expression pattern for transcript ENST00000553192.5.1 with other isoforms. All the above evidence showed that genes with switched isoforms are often functional important in tumorigenesis, but may be ignored from previous studies due to their genes expression may be not significantly differentiated.

**Table 1.** Statistic summary of splicing factor genes with alternative switched isoforms.

| isoform_id | gene_id | gene_name | dIF | q_value |
| --- | --- | --- | --- | --- |
| ENST00000555295.1 | ENSG00000100836.10 | PABPN1 | 0.182 | 1.10E-32 |
| ENST00000459687.5 | ENSG00000100410.7 | PHF5A | 0.172 | 6.07E-18 |
| ENST00000411938.1 | ENSG00000128534.7 | LSM8 | 0.169 | 2.49E-19 |
| ENST00000553192.5 | ENSG00000139343.10 | SNRPF | 0.152 | 6.22E-21 |
| ENST00000297157.7 | ENSG00000164610.8 | RP9 | 0.145 | 2.68E-19 |
| ENST00000491106.1 | ENSG00000060688.12 | SNRNP40 | 0.128 | 4.28E-19 |
| ENST00000560313.2 | ENSG00000090470.14 | PDCD7 | 0.124 | 2.17E-06 |
| ENST00000301785.5 | ENSG00000214753.2 | HNRNPUL2 | 0.116 | 1.19E-28 |
| ENST00000402849.5 | ENSG00000100028.11 | SNRPD3 | 0.113 | 1.67E-11 |
| ENST00000535326.1 | ENSG00000110107.8 | PRPF19 | 0.103 | 1.79E-07 |
| ENST00000597776.1 | ENSG00000130520.10 | LSM4 | 0.102 | 2.31E-34 |
| ENST00000472237.5 | ENSG00000132792.18 | CTNNBL1 | 0.102 | 5.80E-13 |
| ENST00000548994.1 | ENSG00000075188.8 | NUP37 | 0.101 | 1.68E-11 |
| ENST00000564651.5 | ENSG00000102978.12 | POLR2C | 0.1 | 3.01E-14 |
| ENST00000505885.1 | ENSG00000096063.14 | SRPK1 | -0.108 | 1.94E-11 |
| ENST00000404603.5 | ENSG00000100028.11 | SNRPD3 | -0.109 | 2.30E-15 |
| ENST00000540127.1 | ENSG00000214753.2 | HNRNPUL2 | -0.116 | 2.99E-48 |
| ENST00000367208.1 | ENSG00000182004.12 | SNRPE | -0.13 | 1.79E-31 |
| ENST00000527554.2 | ENSG00000100697.14 | DICER1 | -0.139 | 2.98E-21 |
| ENST00000595761.1 | ENSG00000213024.10 | NUP62 | -0.157 | 3.62E-31 |
| ENST00000488937.1 | ENSG00000136875.12 | PRPF4 | -0.159 | 6.94E-12 |
| ENST00000559051.1 | ENSG00000090470.14 | PDCD7 | -0.163 | 9.60E-15 |
| ENST00000216252.3 | ENSG00000100410.7 | PHF5A | -0.216 | 4.30E-21 |

### 3.4 Prediction of AS correlated non-coding RNAs at both transcript and gene level

In order to identify which lncRNAs are associated switched isoforms at the transcript level, we constructed a lncRNA and genes with switched isoforms co-expression network. Different from traditional gene level co-expression network, the connections between lncRNA and genes with multiple splicing isoforms could be 1 vs. 1 or 1 vs. many. The lncRNAs -switched isoforms connections have been summarized in Table S3. Due to space limitation, we only illustrated the relationships between lncRNAs and genes with enriched AS patterns in Fig 4A.

**Figure 4.**
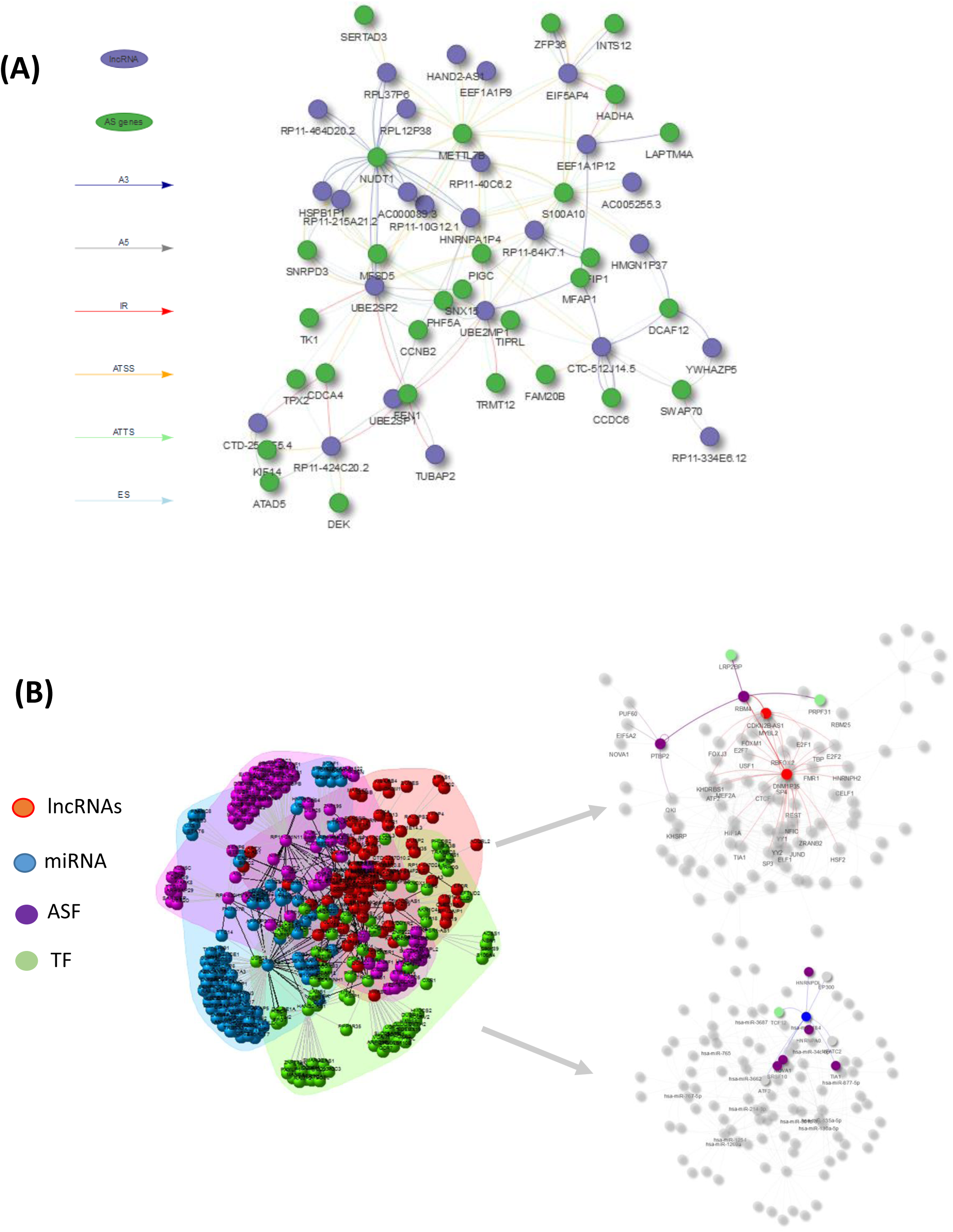
(A) Visulization of lncRNA –AS co-expression network integrated by AS event types (i.e, A3, IR, ES) at the isoform expression level. (B) Illustration of lncRNA-AS comprehive network derived from gene level co-expression network and regulatory network involved with co-effecors miRNA-, TF-, and ASF-interactions.

However, since the lncRNA regulation mechanism involved in AS events is very comprehensive, as the regulation may not directly be reflected from expression abundance, but through physical interaction or DNA/RNA binding. LncRNA could influence genes splicing patterns by inhibiting and activating the expression of alternative splicing factors, or through transcript factors indirectly interact with splicing factors and ultimately cause AS factor targeted gene expression change. Therefore, we constructed a comprehensive gene level network keeping as many TF, AS-regulators, and their targeted genes as possible in AS modulation.

Fig 4B illustrated the HCC lncRNAs -AS network including connections with TFs, ASFs, and miRNAs based on evidence from publicly available resource and gene-level co-expression analysis. Only the lncRNA which directly alters AS genes expression or indirectly alters AS genes through TF, ASF or miRNAs can be included for downstream RWMG analysis. Table S4 provided the prediction of all AS-related genes ranking by RWMG important score.

### 3.5 Computational, Clinical and experimental evaluation for predicted pathogenic lncRNAs involved in AS regulation

As the ROC curve shown in Fig 5A, the averaged area under curve (AUC) value after optimization has been improved from 0.751 to 0.923 based on bootstrapping value. In order to select the best number of top *n* ranked genes that correspond to a good tradeoff between the sensitivity and specificity, we selected the cutoff based on the trend of the changes at which *n* where Δ*TPR* /Δ*FPR* exhibiting a sudden drop (Fig 5B). We can see from the figure, *n*=150 is the best number for gene selection. The top ranked lncRNAs associated with AS functions can be found in Table 2.

**Table 2.** Statistic summary of predicted top-ranked non-coding RNAs associated with AS ranking by RWMG score.

| Gene. Symbol | Ranking | Score | Types |
| --- | --- | --- | --- |
| LINC00675 | 1 | 0.00368174 | lincRNA |
| CTD-2171N6.1 | 2 | 0.002824633 | lincRNA |
| HOTTIP | 3 | 0.002677841 | antisense |
| DNM1P35 | 4 | 0.002483954 | antisense |
| LEF1-AS1 | 5 | 0.002397948 | antisense |
| AP006285.7 | 6 | 0.002123275 | lincRNA |
| WARS2-IT1 | 7 | 0.002081091 | antisense |
| LINC00355 | 8 | 0.002063983 | lincRNA |
| RP11-81H3.2 | 9 | 0.002032482 | lincRNA |
| HOXA11-AS | 10 | 0.002028198 | antisense |
| RP11-261N11.8 | 11 | 0.002005615 | antisense |
| RP3-355L5.4 | 12 | 0.001938399 | antisense |
| RP11-138J23.1 | 13 | 0.001929433 | lincRNA |
| RP11-525K10.3 | 14 | 0.001923733 | antisense |
| RP11-495P10.7 | 15 | 0.001917542 | lincRNA |
| DLX6-AS1 | 16 | 0.00188837 | antisense |
| RP11-356C4.5 | 17 | 0.001861006 | lincRNA |
| CDKN2B-AS1 | 18 | 0.001856714 | antisense |
| RP11-495P10.5 | 19 | 0.001834267 | lincRNA |
| SFTA1P | 20 | 0.001751498 | lincRNA |
| PRSS51 | 21 | 0.001750058 | antisense |
| MALAT1 | 22 | 0.001672339 | lincRNA |
| FEZF1-AS1 | 23 | 0.001669135 | antisense |
| RP4-530I15.9 | 24 | 0.001619806 | antisense |
| RP11-158M2.5 | 25 | 0.001618054 | antisense |
| CTD-2374C24.1 | 26 | 0.001617345 | lincRNA |
| PWRN1 | 27 | 0.001605646 | lincRNA |
| CTC-573N18.1 | 28 | 0.001534221 | lincRNA |
| RP11-284F21.9 | 29 | 0.001527715 | lincRNA |
| RP11-3J1.1 | 30 | 0.001523171 | lincRNA |
| FENDRR | 31 | 0.001509286 | lincRNA |
| hsa-miR-3662 | 1 | 0.003384 | miRNA |
| hsa-miR-760 | 2 | 0.002603 | miRNA |
| hsa-miR-34c-5p | 3 | 0.002485 | miRNA |
| hsa-miR-214-3p | 4 | 0.002232 | miRNA |
| hsa-miR-301b-3p | 5 | 0.002073 | miRNA |
| hsa-miR-3687 | 6 | 0.001947 | miRNA |
| hsa-miR-767-5p | 7 | 0.00189 | miRNA |
| hsa-miR-765 | 8 | 0.00184 | miRNA |
| hsa-miR-1269a | 9 | 0.001744 | miRNA |
| hsa-miR-135a-5p | 10 | 0.001726 | miRNA |
| hsa-miR-1254 | 11 | 0.001669 | miRNA |
| hsa-miR-877-5p | 12 | 0.001662 | miRNA |
| hsa-miR-130a-5p | 13 | 0.001588 | miRNA |
| hsa-miR-184 | 14 | 0.001578 | miRNA |

**Figure 5.**
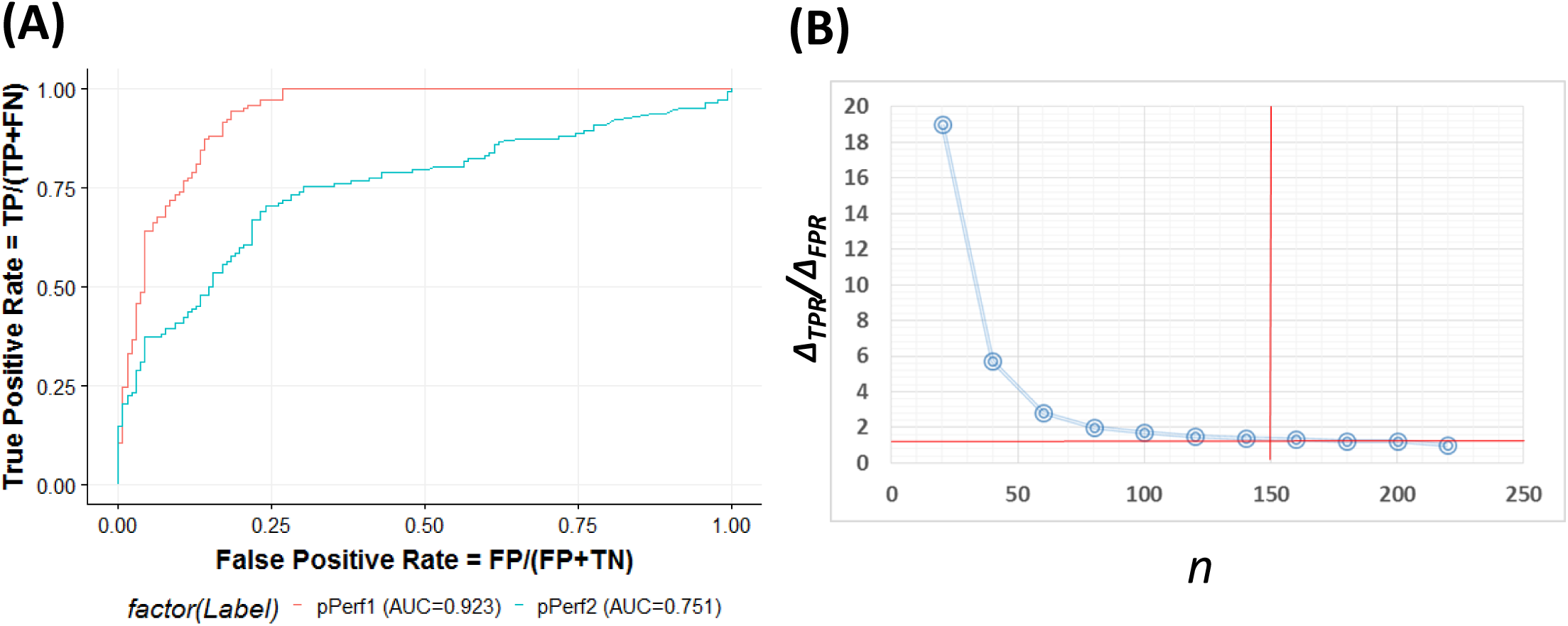
(A) ROC curve for the predictive model evaluation. pPerf1 (AUC=0.923) with the “seed” genes showed a better performance than pPerf2 (AUC=0.751) without the “seed” genes. (2) Trade-off between the sensitivity and specificity with the number of top *n* genes. We can see that the best cutoff is *n*=150, as the ΔTPR/ΔFPR value decreasing very fast in the begining and approaching to smaller changes for n around 150.

Among the top predicted lncRNAs that are involved in AS, we further confirmed their clinical significance. As a result of univariate survival analysis screen, a total of 51 lncRNAs and 24 pseudogenes were found to be associated with HCC overall survival respectively (Table S5). Fig 6 A and B showed the top 10 significant genes based on the Cox proportional regression model. Fig 6 C-D showed the survival curve and distribution of CDKN2B-AS1 and UBE2SP1.

**Figure 6.**
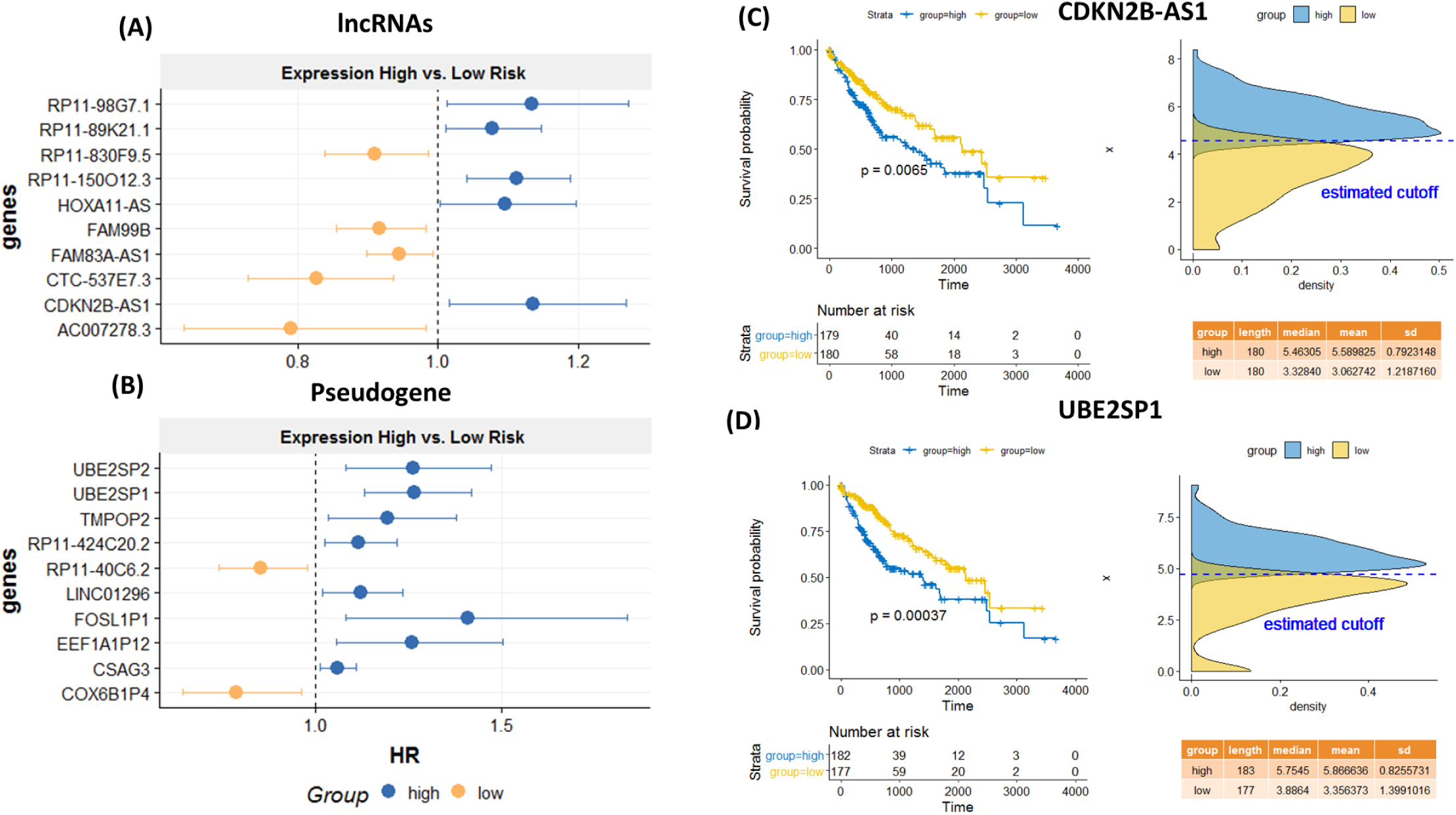
Survival analysis for the identified lncRNA and psudeogenes involved in AS mechanisms. Hazard Ratio plots from Cox regression analysis for top 10 lncRNAs (A) and top 10 pseudogenes (B) associated with overall survival. K-M curves for a lncRNA, CDKN2B-AS1 (C) and a psudogene, UBE2SP1 (D) using the median value as the cutoff. High expression of both genes are significantly associated with poor prognosis.

## 4. Discussion

In the last decade, great studies have investigated the association of splicing isoforms and lncRNAs profiles from deep sequencing. For example, it has been known for a long time that some small nuclear uridine (U)-rich RNAs (snRNPs) who are the core components of the pre-mRNA processing spliceosome can collaborate with some splicing factors which are encoded by heterogeneous nuclear ribonucleoprotein complex subunits (hnRNPs) to fine-tune complex splicing regulations^4^. Impressively, we found a number of core snRNPs isoforms including SNRPE, SNRPD3, SNRPD3, SNRPF, and SNRNP40 are switched, although their expression abundance is not necessary differentially expressed from T/N comparison in HCC development. For example, SNRNP40 catalyzes the removal of introns from pre-messenger RNAs. Similarly, a hnRNP U like protein HNRNPUL2, who is also a scaffold-attachment factor, plays an important role in the formation of a ’transcriptional’ complex by binding to scaffold attachment region and cause chromatin remodeling.

The primary mechanisms involving lncRNAs in AS modulation can be classified in three ways, including: (i) lncRNAs directly influencing isoform expression through activation or suppression mechanism; (ii) lncRNAs forming RNA-RNA duplexes with pre-mRNA molecules and (iii) lncRNAs affecting the target AS genes through indirectly inhibiting or promoting the expression of splicing factors or through transcript factors. However, most previous studies only focus on individual genes and/or isoform switches regulated by lncRNAs. More comprehensive interactions can be detected at isoforms level besides of gene level. Our predictions identified several candidates onco-and tumor suppressor lncRNAs whose somatic alterations associated with AS at both isoform and gene level and of clinical significance in HCC.

In the transcriptional level correlation network, we found the majority of lncRNA isoforms were correlated with more than one AS event, among which some were playing opposite roles in the AS regulations. In addition, we can see that many lncRNAs may partially in competition with the same AS event. For example, the pseudogenes of UBE2S, which are UBE2SP1, UBE2SP2, and UBE2MP1, are significantly correlated with FEN1’s Intron Retention and Alternative 5’ donor sites mechanisms (Fig 4A). The FEN1 gene plays an important role in removing 5’ overhanging flaps and the 5-3 exonuclease activities involved in DNA replication and repair ^47^. While the UBE2S is involved in ubiquitination and subsequent degradation of VHL, resulting in an accumulation of HIF1A ^48^, however, the reason why its pseudogenes are associated with FEN1 is not clear. Maybe the next step for experimental validation for a deep understanding of the mechanisms. Taken together, these results confirmed that these identified lncRNAs need to be better investigated in the future. Our results provided a better resolution of AS correlated lncRNAs at the isoform level.

AS events are mainly regulated by splicing factors, which bind to pre-mRNAs and influence exon selection and splicing site choice. Moreover, transcriptional factors will activate or suppress the expression of ASF. Importantly, we found ASF may have switched isoforms. A switched ASF RP9, which can be bound by the proto-oncogene PIM1 product, a serine/threonine protein kinase, also can cause its target PIM1 switched. Although TFs were usually thought for a long time to encode a single protein that changes the expression of their target genes, more and more TFs, are now found to be alternatively spliced ^49^. Here, we also found a group of TFs in the ETS family (E26 transformation-specific), which are ETS1, ETS2, ETV3, ELF4, are switched together. These ETS genes have been confirmed to be associated with cancer through gene fusion ^50^ and are involved in a wide variety of regulatory functions such as cell migration, proliferation, and cancer progression ^51 52^. Interestingly, the ETS1 target splicing factor QKI, and ETV3 target splicing factor CELF1. More interestingly, lncRNA FAM99B is predicted to be associated with these ETS family genes and their low expression are associated with HCC patients’ poor prognosis.

CDKN2B-AS1, also known as ANRIL, its association with HCC has been reported in several studies^53-55^. CDKN2B-AS1 has both linear and circular isoforms and their functions are different. For example, its linear isoform can regulate the c-myc-enhancer binding factor RBMS1 ^56^, while its circular isoform is confirmed to be an important AS regulator that causing skipped exons ^57^ mainly found in cardiovascular disease ^58 59^. However, this is the first time we found it can activate alternative splicing genes in liver cancer. A potential explanation could be it is functionally related to lipid metabolism and a majority of liver cancer due to lipid disorder. In addition, the prognostic value of CDKN2B-AS1 has been revealed in our project. However, how exactly CDKN2B-AS1 controls these genes’ splicing is not clear. Further experimental validation can be planed. We identified HAND2-AS1 gene showed consistent alternative splicing pattern at the start sites and termination site for METTL7B at isoforms level. METTL7B is a membrane-associated protein that resides on hepatic lipid Droplets. An explanation is that HAND2-AS1 activate the METTL7B spliced isoform lipid disordered and is associated with HCC, which didn’t report before. Gene-level RWMG network analysis further reveals that both CDKN2B-AS1 and HAND2-AS1 can influence AS either through TFs and ASFs. For example, HAND2-AS1 TFs (i.e. ETS1, SP1, E2F7), or ASFs (i.e. SRSF7, SFRP1, HNRNPK); CDKN2B-AS1 associated TFs (SP4, E2F7) and ASF (SRSF1, SRSF2).

In this project, we extended this algorithm to multiplex and heterogeneous networks. The walk can explore different layers of the epigenetic regulatory network, expression correlation network, and protein interaction network. A recent Nature Review paper by Sharan et al., also suggested that the “network-propagation” method is a “powerful” and “accurate” refined approach in the network, since it is capable of dealing with “noisy” and “incomplete” observations by simultaneously considering all possible paths among vertices. Analyzing these heterogeneous data together will significantly improve the prediction accuracy. By using this gene-ranking strategy, potentially spurious predictions (false positives) that are supported by a single (shortest) path are down-weighted, and true high ranked genes that are potentially missed, even though they are well connected to the prior list (false negatives), are promoted.

To our best knowledge, this is the first attempt to predict lncRNAs regulations on AS using a rigorous, multi-graphic approach by integrating such large scale and complex networks. Of interest for potentially limiting the accuracy of random walk and network propagation methods are an incomplete collection of known lncRNAs, especially pseudogenes, used to supervise prediction of new candidates. As such, we addressed several unique challenges associated with these datasets’ complexity in each step. For example, in the data preprocessing steps, we have carefully addressed the challenges by collecting as many as experimental verified and predicted lncRNAs that are taking account of AS; In our statistical modeling steps, we carefully addressed the robustness of complex data integration, especially for non-informative or noisy datasets. Also, we investigated several random walk strategies by trying different groups of vertices such as lncRNAs, ASFs and TFs as a start point to optimize models.

However, the lncRNA regulatory mechanism is very complicated, as its mechanism differs with different stages, such as the pre-mRNA or post-mRNA stage. Therefore, the main limitation of this project is we are not able to consider several other comprehensive mechanisms at different stages, such as recognition of the splicing site can be modulated by cis-regulatory sequences, known as splicing enhancers or silencers, which contribute to the generation of two or more alternatively-spliced mRNAs from the same pre-mRNA. Also, lncRNA determine AS patterns through chromatin remodeling mechanism and shape the 3D genome organization. Our next step will focus on interpreting these mechanisms at different stages.

## 5. Conclusion

We performed a large-scale RNAseq analysis to identify alternative splicing isoforms and their biological consequences in liver cancer. To predict which lncRNAs are associated with splicing mechanisms in HCC, we developed a RWMG model to integrate multi-layers heterogeneous networks including epigenetic regulatory network, transcriptional level co-expression network, and PPI network with the evidence that lncRNAs in the correlation between effectors (miRNAs, TFs, or ASFs) and their associated splicing genes. Our project is the first time using the network-based computational method to genome-wisely predict AS-related lncRNAs in HCC, which showed a good prediction performance (AUC=0.923) and clinical significance.

## Acknowledgements

N/A

## Author contributions

JW contributed to the analysis and interpretation of data and reviewing of the manuscript. YC and KX contributed to the machine learning predictive model design. YM contributed to the project design and interpretation of biological meanings. YZ led the project, provided guidance and prepared the manuscript. All authors read and approved the manuscript

## Supplementary files

**Figure S1**. (A) Illustrations of overall project design, and (B) explaination of biological mechanisms.

**Table S1**. Statistic summaries for significantly differential expression genes between tumor and normal comparison for lncRNAs and pseudogenes.

**Table S2.** Statistic summaries and functional analysis for significantly switched isoforms between tumor and normal comparison for lncRNA and protein-coding genes, as well as prediction of splicing event patterns for switched genes. Gene ontology and KEGG pathway enrichment analysis are performed for upregulated isoforms (gain) and downregulated isoforms (loss) respectively.

**Table S3.** Predicted lncRNA interaction pairs at the transcriptional co-expression level.

**Table S4.** Statistic summary of AS-associated lncRNAs and PCGs ranking by RWMG predictive score.

**Table S5.** Statistic summary of significant lncRNAs and psudogenes associated with overall survival.

## References

Climente-Gonzalez H, Porta-Pardo E, Godzik A, Eyras E. The functional impact of alternative splicing in cancer. Cell reports 2017; 20(9): 2215–26.

Eskens FA, Ramos FJ, Burger H, et al. Phase I, pharmacokinetic and pharmacodynamic study of the first-in-class spliceosome inhibitor E7107 in patients with advanced solid tumors. Clinical Cancer Research 2013: clincanres. 0485.2013.

Zhang L, Liu X, Zhang X, Chen R. Identification of important long non-coding RNAs and highly recurrent aberrant alternative splicing events in hepatocellular carcinoma through integrative analysis of multiple RNA-Seq datasets. Molecular Genetics and Genomics 2016; 291(3): 1035–51.

Romero-Barrios N, Legascue MF, Benhamed M, Ariel F, Crespi M. Splicing regulation by long noncoding RNAs. Nucleic acids research 2018; 46(5): 2169–84.

Wang J, Su L, Chen X, et al. MALAT1 promotes cell proliferation in gastric cancer by recruiting SF2/ASF. Biomedicine & Pharmacotherapy 2014; 68(5): 557–64.

West JA, Davis CP, Sunwoo H, et al. The long noncoding RNAs NEAT1 and MALAT1 bind active chromatin sites. Molecular cell 2014; 55(5): 791–802.

Ji Q, Zhang L, Liu X, et al. Long non-coding RNA MALAT1 promotes tumour growth and metastasis in colorectal cancer through binding to SFPQ and releasing oncogene PTBP2 from SFPQ/PTBP2 complex. British journal of cancer 2014; 111(4): 736.

Kong J, Sun W, Li C, et al. Long non-coding RNA LINC01133 inhibits epithelial–mesenchymal transition and metastasis in colorectal cancer by interacting with SRSF6. Cancer letters 2016; 380(2): 476–84.

Zang C, Nie F-q, Wang Q, et al. Long non-coding RNA LINC01133 represses KLF2, P21 and E-cadherin transcription through binding with EZH2, LSD1 in non small cell lung cancer. Oncotarget 2016; 7(10): 11696.

David CJ, Chen M, Assanah M, Canoll P, Manley JL. HnRNP proteins controlled by c-Myc deregulate pyruvate kinase mRNA splicing in cancer. Nature 2010; 463(7279): 364.

Koh CM, Bezzi M, Low DH, et al. MYC regulates the core pre-mRNA splicing machinery as an essential step in lymphomagenesis. Nature 2015; 523(7558): 96.

Vivian J, Rao AA, Nothaft FA, et al. Toil enables reproducible, open source, big biomedical data analyses. Nature Biotechnology 2017; 35(4): 314–6.

Harrow J, Frankish A, Gonzalez JM, et al. GENCODE: the reference human genome annotation for The ENCODE Project. Genome research 2012; 22(9): 1760–74.

Robinson MD, Oshlack A. A scaling normalization method for differential expression analysis of RNA-seq data. Genome biology 2010; 11(3): R25.

Smyth GK. Limma: linear models for microarray data. Bioinformatics and computational biology solutions using R and Bioconductor: Springer; 2005: 397–420.

Wang J, Zhou Y, Fei X, Chen X, Chen Y. Biostatistics mining associated method identifies AKR1B10 enhancing hepatocellular carcinoma cell growth and degenerated by miR-383-5p. Scientific reports 2018; 8.

Vitting-Seerup K, Sandelin A. IsoformSwitchAnalyzeR: Analysis of changes in genome-wide patterns of alternative splicing and its functional consequences. bioRxiv 2018: 399642.

Vitting-Seerup K, Sandelin A. The landscape of isoform switches in human cancers. Molecular Cancer Research 2017.

Wang L, Park HJ, Dasari S, Wang S, Kocher J-P, Li W. CPAT: Coding-Potential Assessment Tool using an alignment-free logistic regression model. Nucleic acids research 2013; 41(6): e74–e.

Finn RD, Coggill P, Eberhardt RY, et al. The Pfam protein families database: towards a more sustainable future. Nucleic acids research 2015; 44(D1): D279–D85.

Potter SC, Luciani A, Eddy SR, Park Y, Lopez R, Finn RD. HMMER web server: 2018 update. Nucleic acids research 2018.

Cuccurese M, Russo G, Russo A, Pietropaolo C. Alternative splicing and nonsense-mediated mRNA decay regulate mammalian ribosomal gene expression. Nucleic acids research 2005; 33(18): 5965– 77.

Weischenfeldt J, Waage J, Tian G, et al. Mammalian tissues defective in nonsense-mediated mRNA decay display highly aberrant splicing patterns. Genome biology 2012; 13(5): R35.

Vitting-Seerup K, Porse BT, Sandelin A, Waage J. spliceR: an R package for classification of alternative splicing and prediction of coding potential from RNA-seq data. BMC bioinformatics 2014; 15(1): 81.

Chiu H-S, Somvanshi S, Patel E, et al. Pan-Cancer analysis of lncRNA regulation supports their targeting of cancer genes in each tumor context. Cell reports 2018; 23(1): 297.

Dweep H, Gretz N. miRWalk2. 0: a comprehensive atlas of microRNA-target interactions. Nature methods 2015; 12(8): 697.

Li J-H, Liu S, Zhou H, Qu L-H, Yang J-H. starBase v2. 0: decoding miRNA-ceRNA, miRNA-ncRNA and protein–RNA interaction networks from large-scale CLIP-Seq data. Nucleic acids research 2013: gkt1248.

Chen G, Wang Z, Wang D, et al. LncRNADisease: a database for long-non-coding RNA-associated diseases. Nucleic acids research 2013; 41(D1): D983–D6.

Bao Z, Yang Z, Huang Z, Zhou Y, Cui Q, Dong D. LncRNADisease 2.0: an updated database of long non-coding RNA-associated diseases. Nucleic acids research 2018.

Consortium EP. The ENCODE (ENCyclopedia of DNA elements) project. Science 2004; 306(5696): 636–40.

Bovolenta LA, Acencio ML, Lemke N. HTRIdb: an open-access database for experimentally verified human transcriptional regulation interactions. BMC genomics 2012; 13(1): 405.

Whitfield TW, Wang J, Collins PJ, et al. Functional analysis of transcription factor binding sites in human promoters. Genome biology 2012; 13(9): R50.

Matys V, Kel-Margoulis OV, Fricke E, et al. TRANSFAC® and its module TRANSCompel®: transcriptional gene regulation in eukaryotes. Nucleic acids research 2006; 34(suppl_1): D108–D10.

Belinky F, Nativ N, Stelzer G, et al. PathCards: multi-source consolidation of human biological pathways. Database 2015; 2015.

Kanehisa M, Goto S. KEGG: kyoto encyclopedia of genes and genomes. Nucleic acids research 2000; 28(1): 27–30.

Geer LY, Marchler-Bauer A, Geer RC, et al. The NCBI biosystems database. Nucleic acids research 2009; 38(suppl_1): D492–D6.

Croft D, O’kelly G, Wu G, et al. Reactome: a database of reactions, pathways and biological processes. Nucleic acids research 2010; 39(suppl_1): D691–D7.

Giulietti M, Piva F, D’Antonio M, et al. SpliceAid-F: a database of human splicing factors and their RNA-binding sites. Nucleic acids research 2012: gks997.

Paz I, Akerman M, Dror I, Kosti I, Mandel-Gutfreund Y. SFmap: a web server for motif analysis and prediction of splicing factor binding sites. Nucleic acids research 2010; 38(suppl_2): W281–W5.

Franceschini A, Szklarczyk D, Frankild S, et al. STRING v9. 1: protein-protein interaction networks, with increased coverage and integration. Nucleic acids research 2012; 41(D1): D808–D15.

Ma C, Chen Y, Wilkins D, Chen X, Zhang J. An unsupervised learning approach to find ovarian cancer genes through integration of biological data. BMC genomics 2015; 16(9): S3.

Fang H, Gough J. Thednet’approach promotes emerging research on cancer patient survival. Genome medicine 2014; 6(8): 64.

Csardi G, Nepusz T. The igraph software package for complex network research. InterJournal, Complex Systems 2006; 1695(5): 1–9.

Guerrieri F. Long non-coding RNAs era in liver cancer. World journal of hepatology 2015; 7(16): 1971.

Kunej T, Obsteter J, Pogacar Z, Horvat S, Calin GA. The decalog of long non-coding RNA involvement in cancer diagnosis and monitoring. Critical reviews in clinical laboratory sciences 2014; 51(6): 344–57.

Quagliata L, Matter MS, Piscuoglio S, et al. Long noncoding RNA HOTTIP/HOXA13 expression is associated with disease progression and predicts outcome in hepatocellular carcinoma patients. Hepatology 2014; 59(3): 911–23.

Wang J, Cao P, Qi Y-Y, et al. The relationship between cell apoptosis dysfunction and FEN1 E160D mutation in lupus nephritis patients. Autoimmunity 2017; 50(8): 476–80.

Jung C-R, Hwang K-S, Yoo J, et al. E2-EPF UCP targets pVHL for degradation and associates with tumor growth and metastasis. Nature medicine 2006; 12(7): 809.

Marcel V, Hainaut P. p53 isoforms-a conspiracy to kidnap p53 tumor suppressor activity? Cellular and Molecular Life Sciences 2009; 66(3): 391.

Tomlins SA, Rhodes DR, Perner S, et al. Recurrent fusion of TMPRSS2 and ETS transcription factor genes in prostate cancer. science 2005; 310(5748): 644–8.

Lee GM, Donaldson LW, Pufall MA, et al. The structural and dynamic basis of Ets-1 DNA binding autoinhibition. Journal of Biological Chemistry 2005; 280(8): 7088–99.

Sharrocks AD. The ETS-domain transcription factor family. Nature reviews Molecular cell biology 2001; 2(11): 827.

Ma J, Li T, Han X, Yuan H. Knockdown of LncRNA ANRIL suppresses cell proliferation, metastasis, and invasion via regulating miR-122-5p expression in hepatocellular carcinoma. Journal of cancer research and clinical oncology 2018; 144(2): 205–14.

Hua L, Wang C-Y, Yao K-H, Chen J-T, Zhang J-J, Ma W-L. High expression of long non-coding RNA ANRIL is associated with poor prognosis in hepatocellular carcinoma. International journal of clinical and experimental pathology 2015; 8(3): 3076.

Chen W-m, Qi F-z, Xia R, et al. Long non-coding RNA ANRIL is upregulated in hepatocellular carcinoma and regulates cell proliferation by epigenetic silencing of KLF2. Journal of hematology & oncology 2015; 8(1): 57.

Hubberten M, Bochenek G, Chen H, et al. Linear isoforms of the long noncoding RNA CDKN2B-AS1 regulate the c-myc-enhancer binding factor RBMS1. European Journal of Human Genetics 2018: 1.

Holdt LM, Stahringer A, Sass K, et al. Circular non-coding RNA ANRIL modulates ribosomal RNA maturation and atherosclerosis in humans. Nature communications 2016; 7: 12429.

Sarkar D, Oghabian A, Bodiyabadu PK, et al. Multiple isoforms of ANRIL in melanoma cells: structural complexity suggests variations in processing. International journal of molecular sciences 2017; 18(7): 1378.

Burd CE, Jeck WR, Liu Y, Sanoff HK, Wang Z, Sharpless NE. Expression of linear and novel circular forms of an INK4/ARF-associated non-coding RNA correlates with atherosclerosis risk. PLoS genetics 2010; 6(12): e1001233.

