## Supplementary figures and images for "Comprehensive network analysis reveals alternative splicing-related lncRNAs in hepatocellular carcinoma"

### Figure S1.pdf

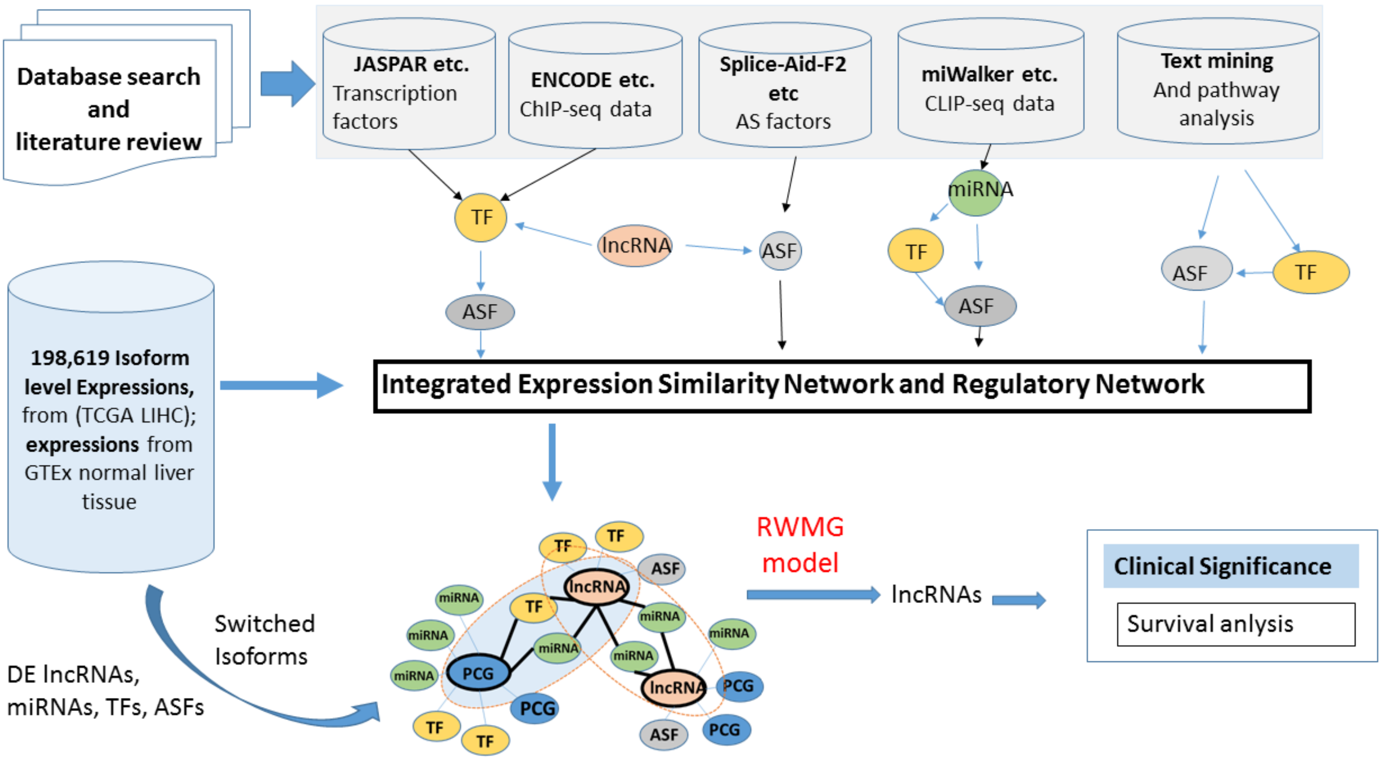

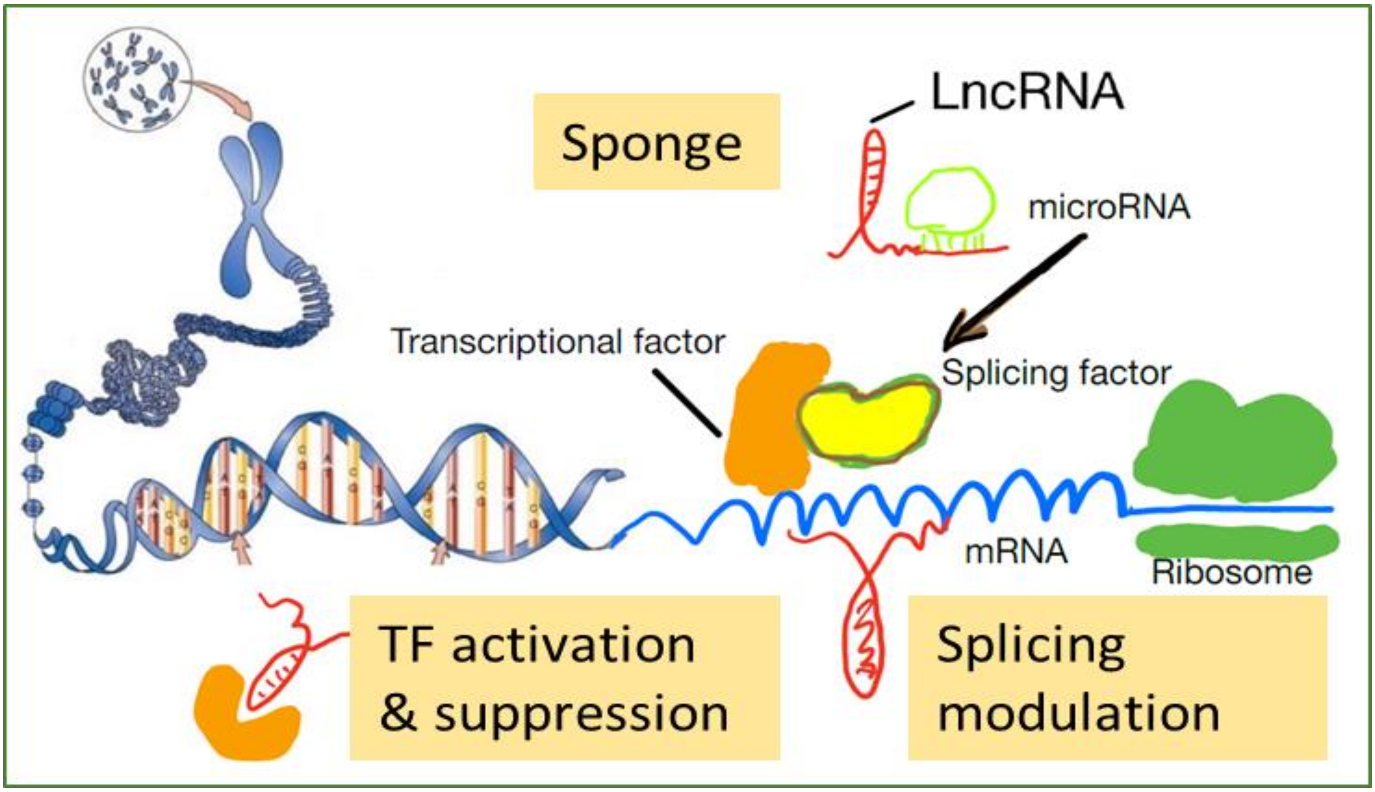
